## Supporting Information for "Origin of Conformational Dynamics in a Globular Protein"

**This file includes:**

Supplementary Figures 1–13
Supplementary Tables 1–2
Supplementary References

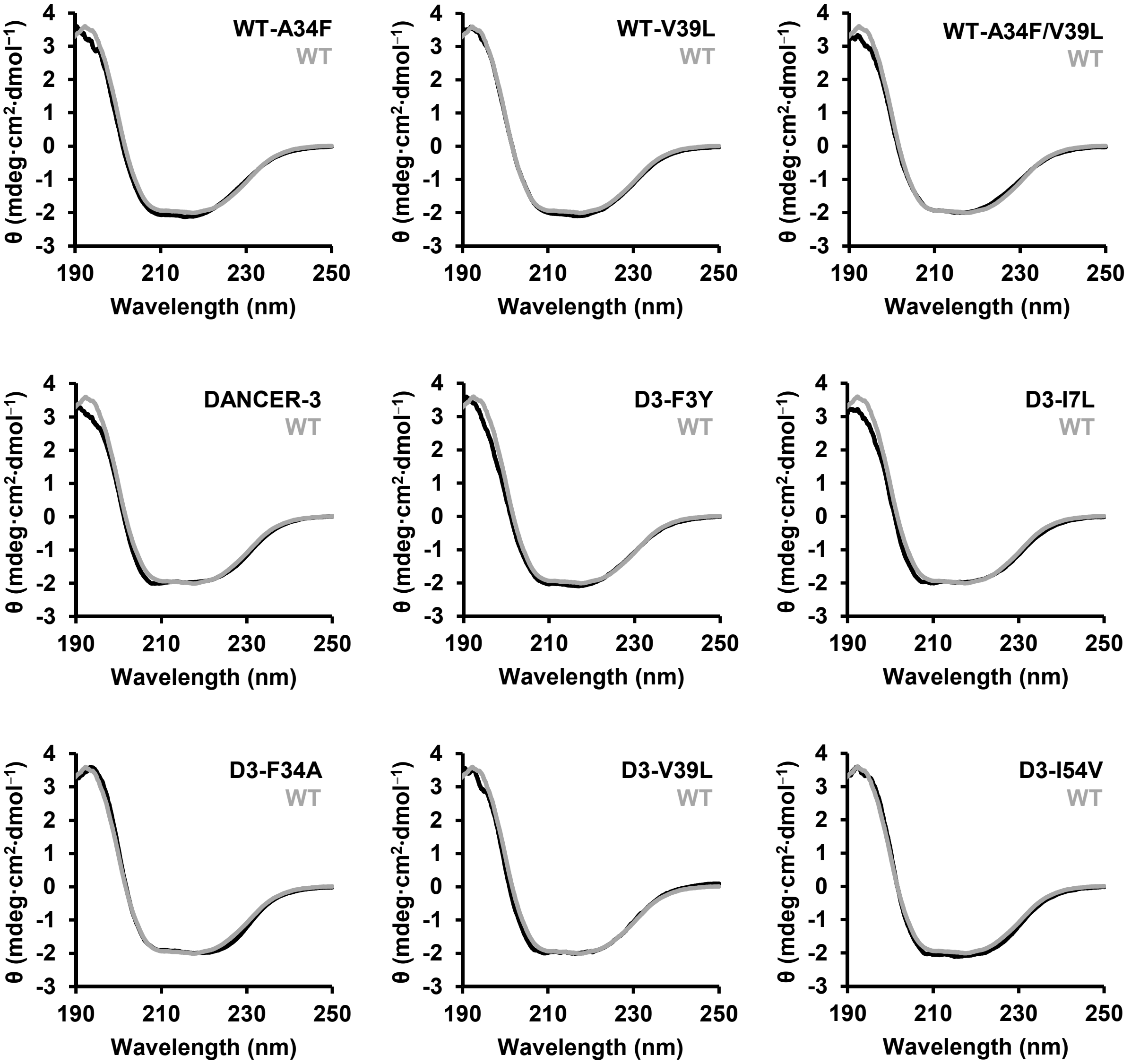


**Supplementary Figure 1. Gβ1 variants adopt the native fold**. Far-UV circular dichroism spectra of Gβ1 variants (black) superimposed on the spectrum of the wild-type (WT) protein (grey).

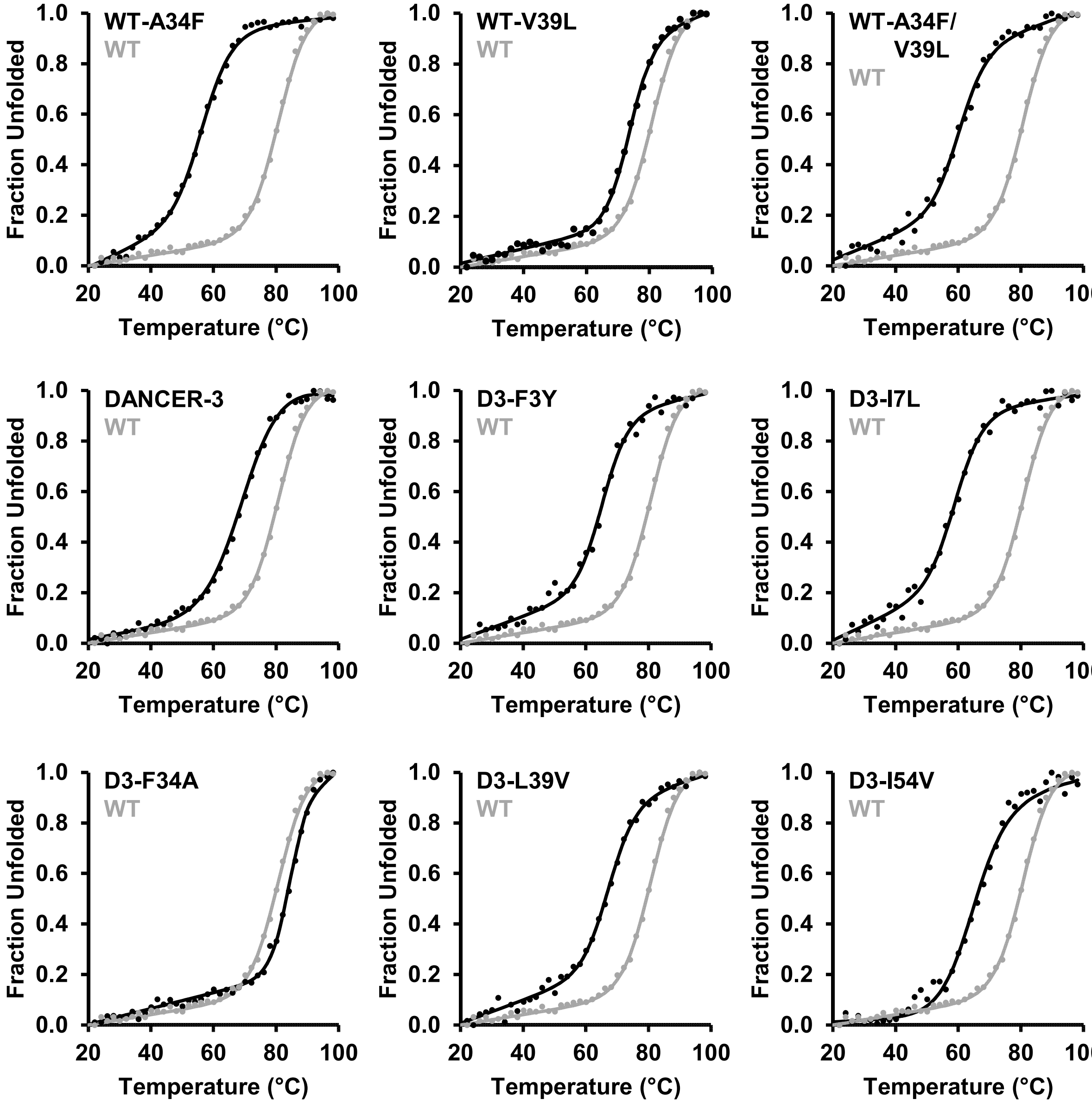


**Supplementary Figure 2. Thermal denaturation of Gβ1 variants.** Thermal denaturation monitored by circular dichroism at 208 nm indicates that all Gβ1 variants are stable at room temperature (n = 1). Data was fit to a two-state unfolding model.[35] Data from the thermal denaturation of wild-type (WT) Gβ1 is shown in grey for comparison.

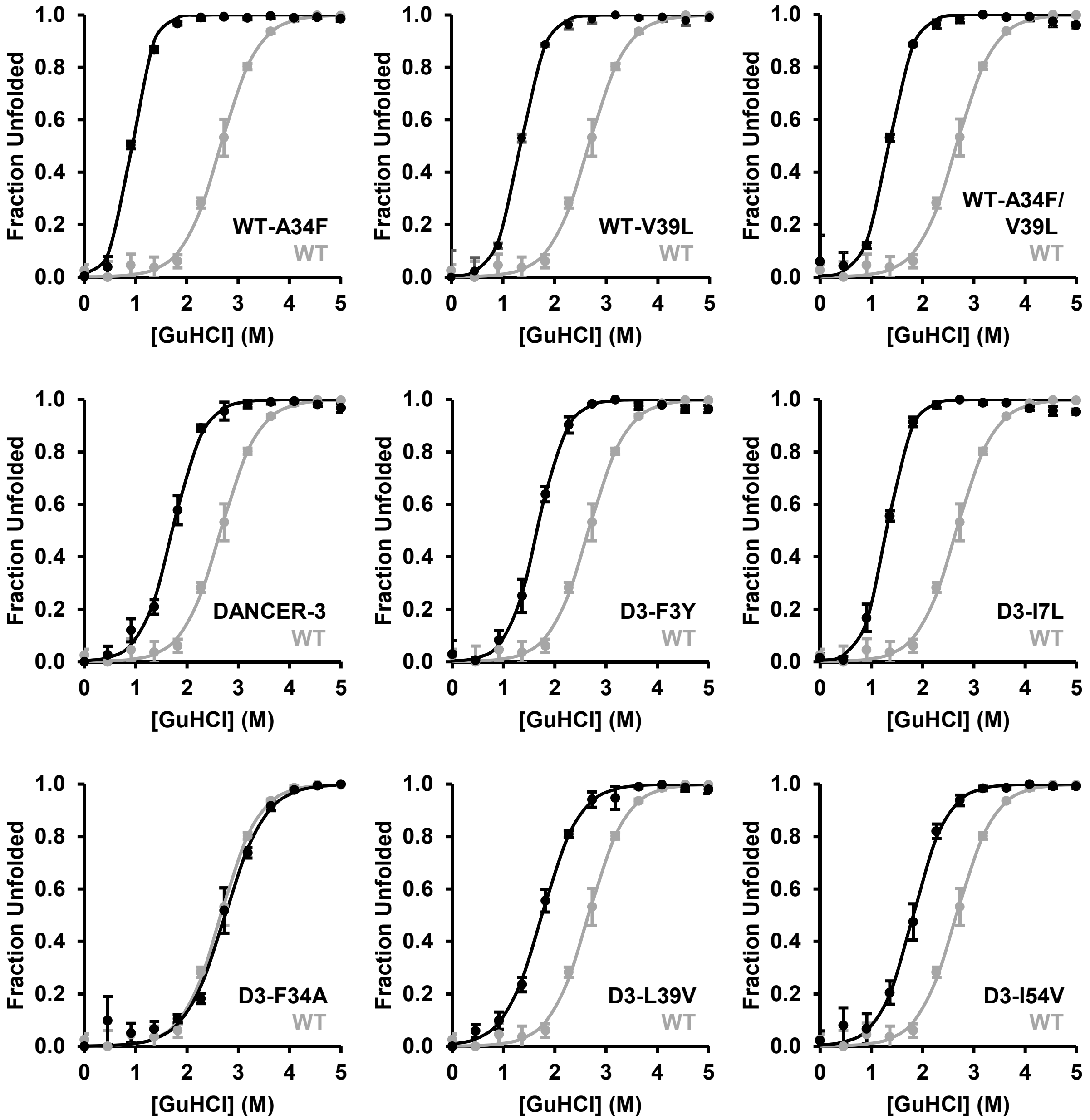


**Supplementary Figure 3. Chemical denaturation of Gβ1 variants.** Chemical denaturation using guanidium chloride (GuHCl) demonstrates that all proteins unfold according to a two-state model (n = 3). Fraction unfolded values were calculated by monitoring integrated tryptophan fluorescence ($\lambda_{excitation}$ = 280 nm, $\lambda_{emission}$ = 310–450 nm) relative to fluorescence at 0 M GuHCl. All experiments were performed in triplicate, and the average and standard deviation reported for each data point. The wild-type (WT) denaturation curve is shown in grey for comparison.

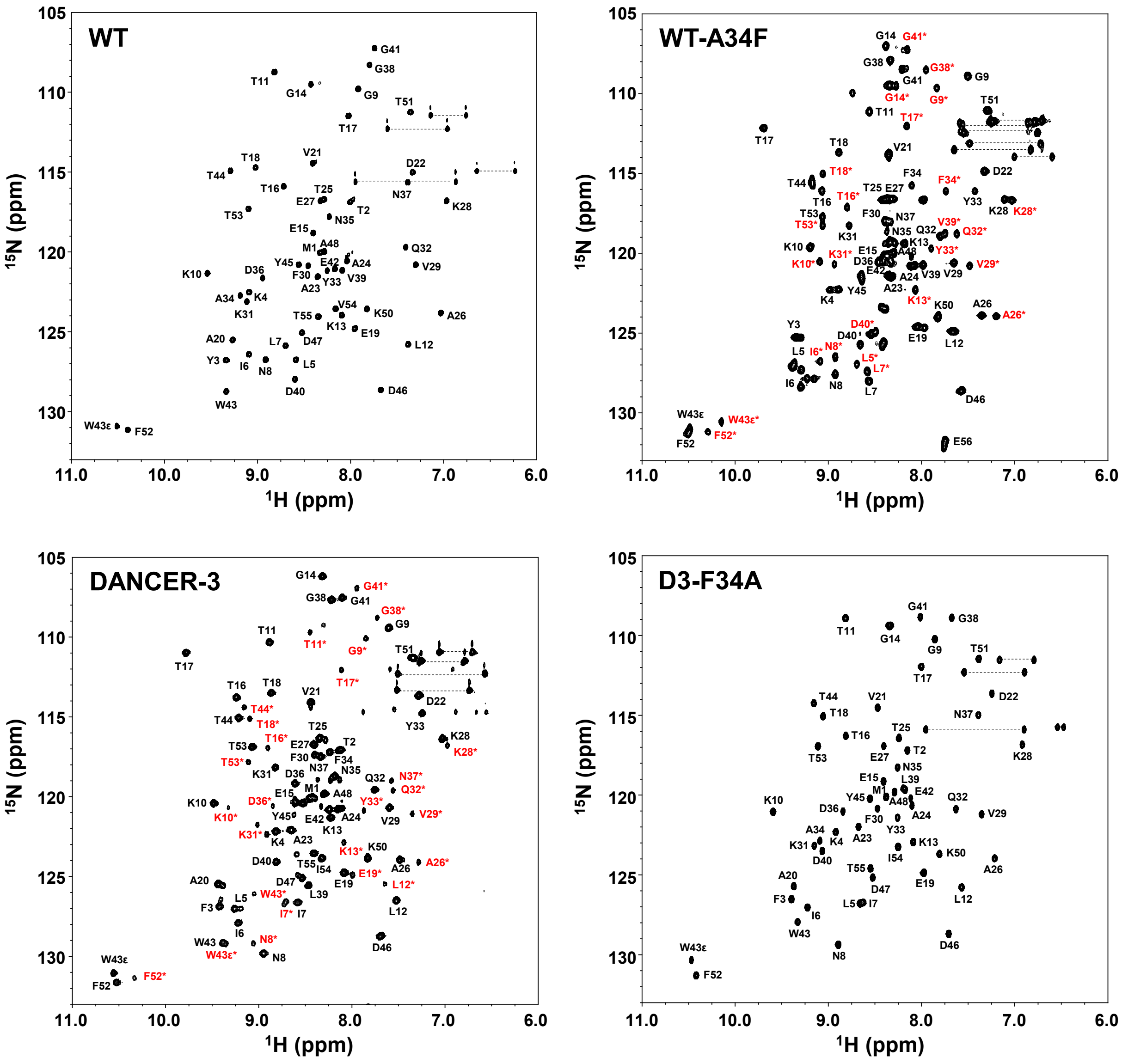


**Supplementary Figure 4. Assigned $^{1}$H-$^{15}$N HSQC spectra of selected Gβ1 variants.** Peaks corresponding to the minor state are indicated with red assignments. Side-chain amide resonances from asparagine and glutamine residues are connected by horizontal lines. WT-A34F assignments were obtained from Jee J., *et al*. (2008).

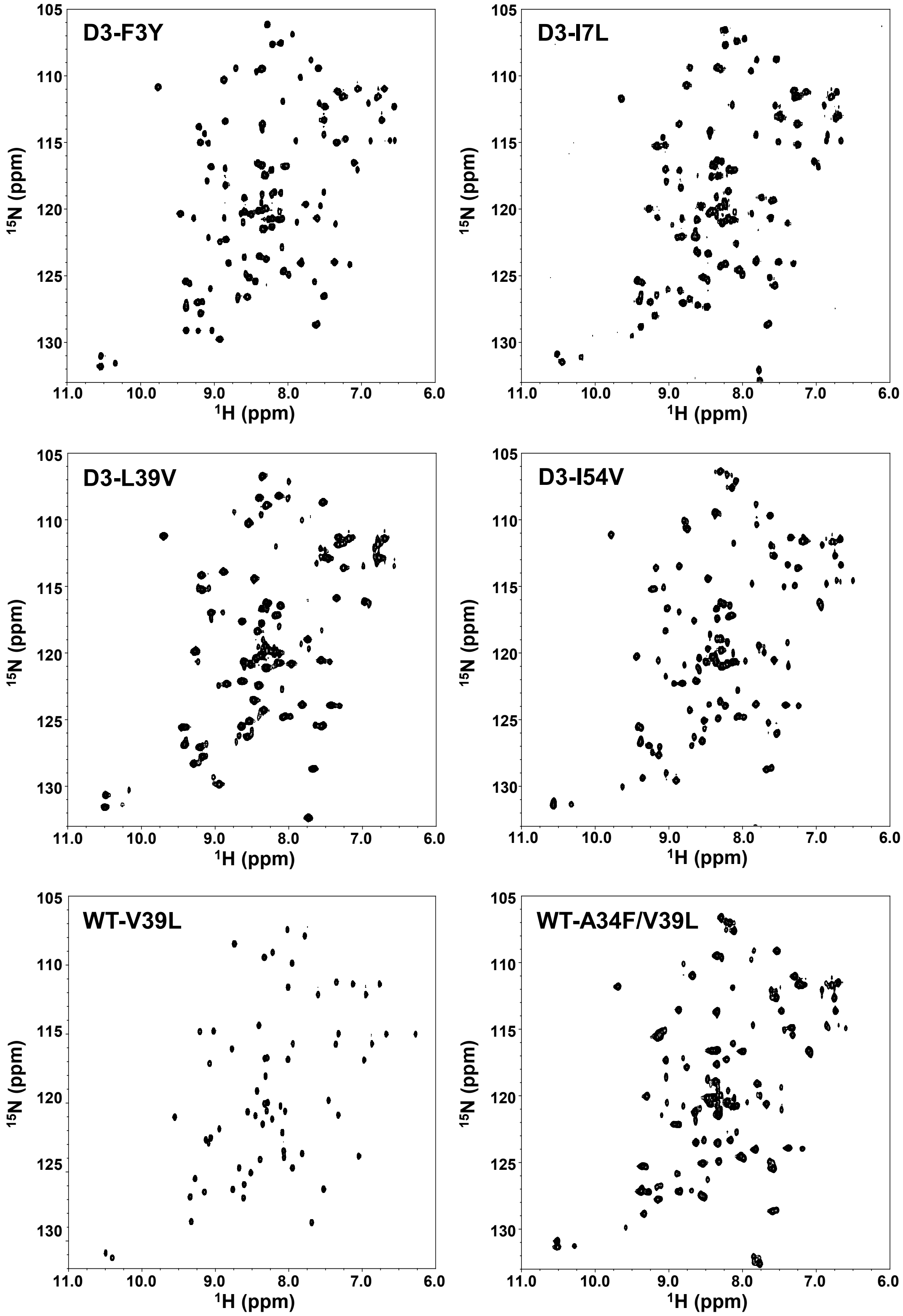


**Supplementary Figure 5. Unassigned $^{1}H$-$^{15}N$ HSQC spectra of selected Gβ1 variants.**

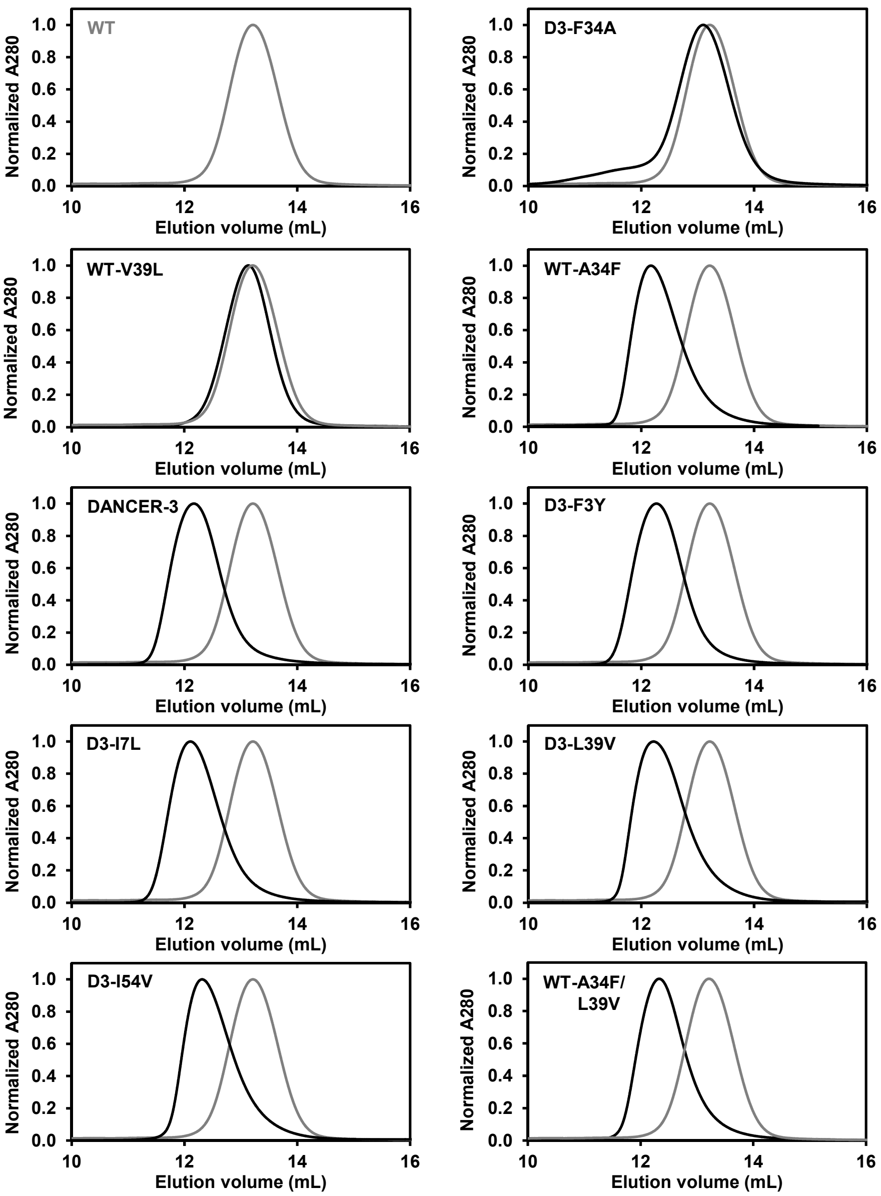


**Supplementary Figure 6. Analytical size-exclusion chromatograms of Gβ1 variants.** Chromatograms monitored by absorption at 280 nm using an AKTA Superdex 75 10/300 column show a shift of ~1 mL in elution volume for dimeric Gβ1 variants relative to the monomeric wild type (WT), shown in grey for comparison.

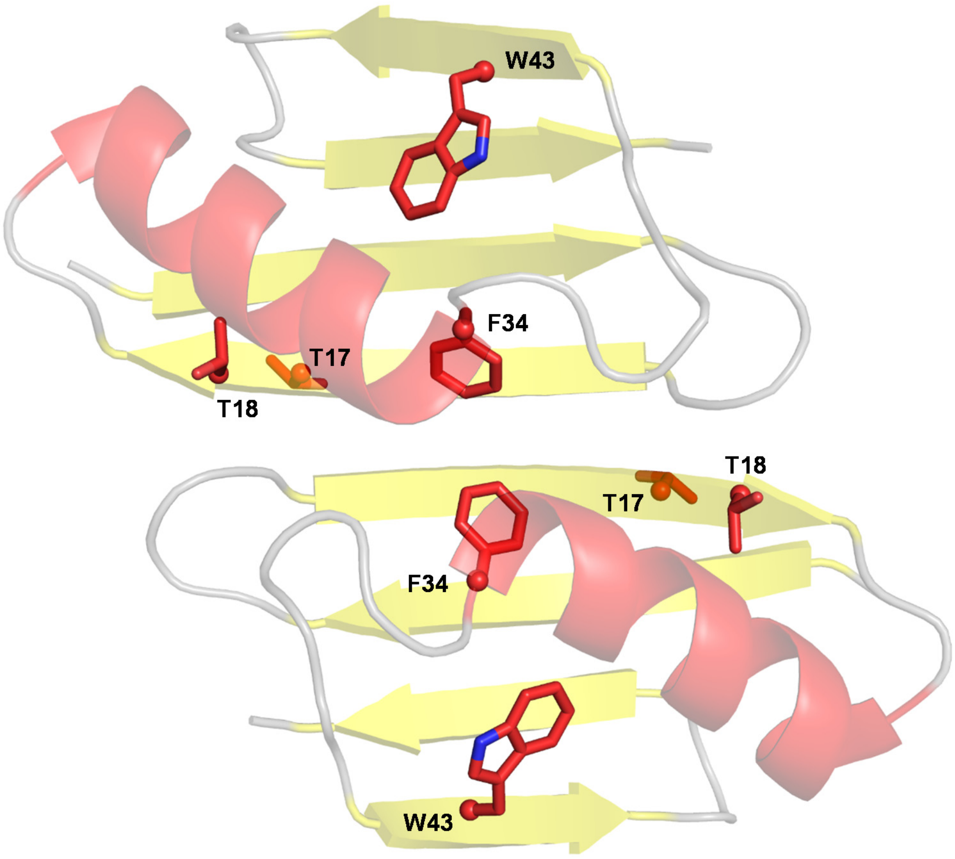


**Supplementary Figure 7. Structure of the WT-A34F dimer.** Solution NMR structure of WT-A34F (PDB ID: 2RMM).[31] F34 is responsible for inducing dimerization, while T17 and T18 act as reporters of dimerization in NMR.

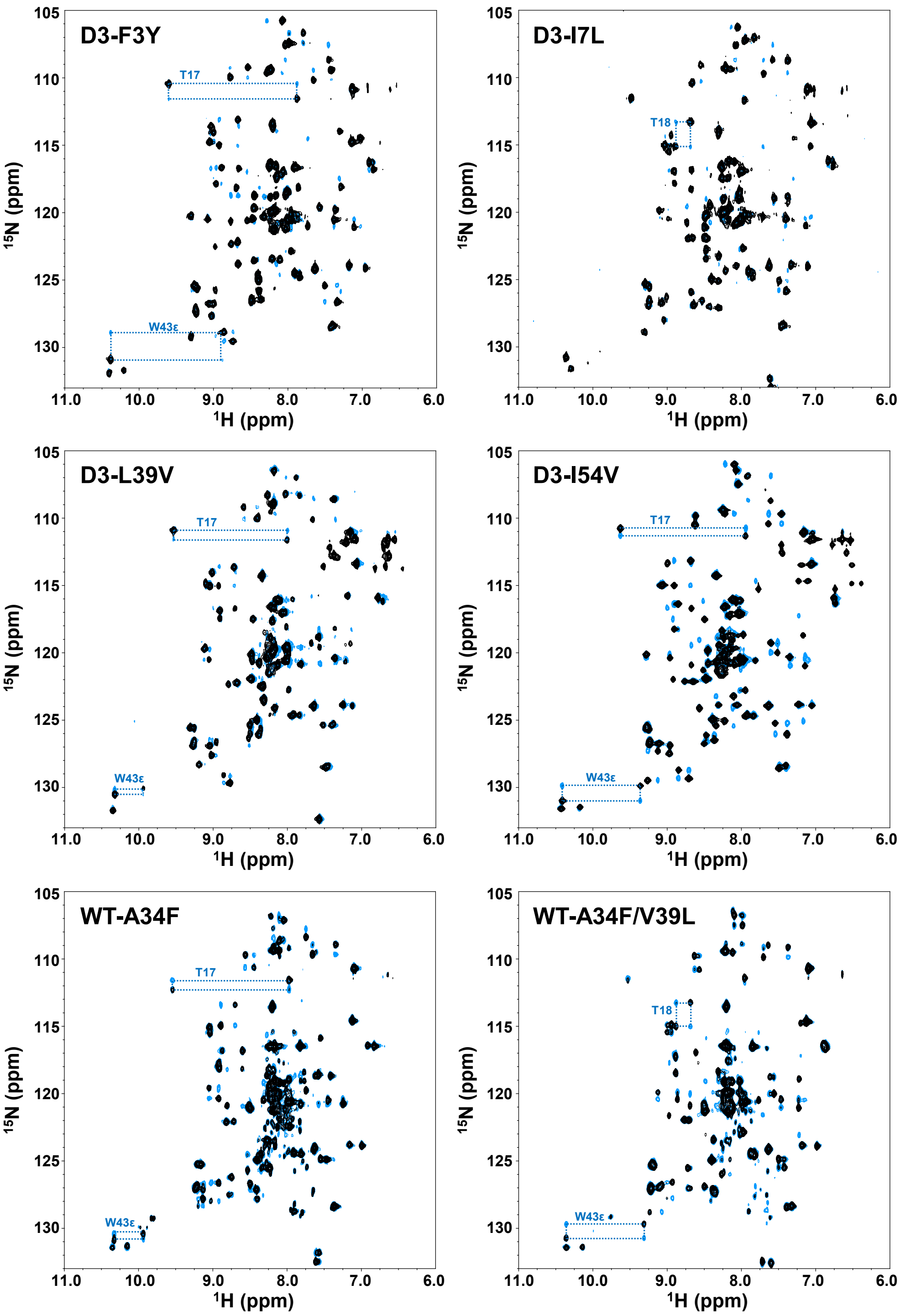


**Supplementary Figure 8. $^1$H-$^{15}$N HSQC ZZ-Exchange spectra of selected Gβ1 variants.** Representative ZZ-exchange spectra are shown in blue and overlaid with $^1$H-$^{15}$N HSQCs in black to highlight the presence of exchange peaks. Dotted lines connect exchange crosspeaks with peaks from the major and minor conformational states used in the analysis of exchange kinetics for each respective variant.

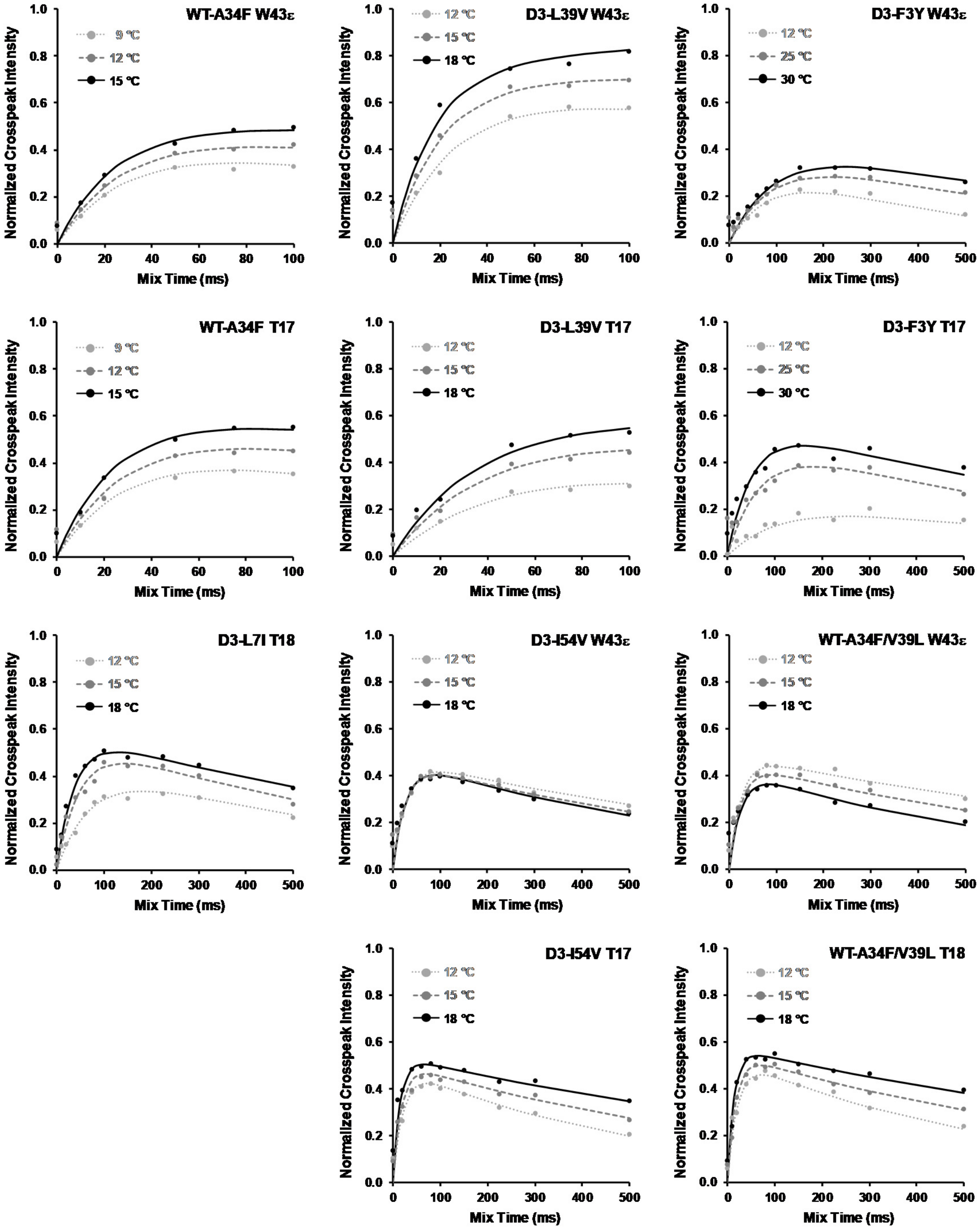


**Supplementary Figure 9. Effect of temperature on exchange rates.** $^{1}$H-$^{15}$N HSQC ZZ-exchange crosspeak intensity curves for the W43ε minor to major state transition are shown for three temperatures per variant and were fitted using a four-term exchange and relaxation model.[38]

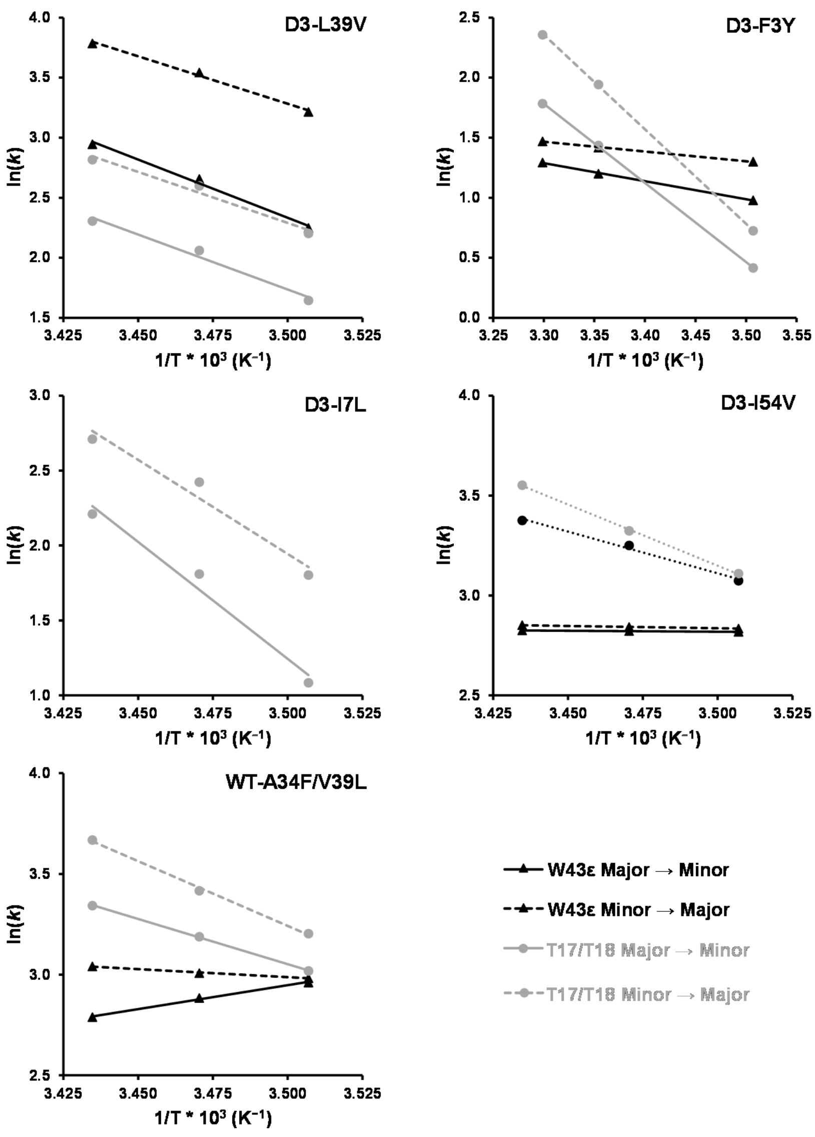


**Supplementary Figure 10. Arrhenius plots of selected Gβ1 variants.** Arrhenius plots demonstrate Arrhenius behavior for both W43ε and T17 exchange profiles in D3-L39V, as shown by similar activation energies for all four transitions. D3-F3Y, D3-I54V, and WT-A34F/V39L however demonstrate non-Arrhenius behavior where one or both W43ε transitions display a non-physical negative or very weak activation energy indicating that they do not follow a simple two-state Arrhenius model, as opposed to the normal Arrhenius behavior demonstrated by T17. W43ε minor state peaks and crosspeaks could not be accurately quantified for D3-I7L due to their low intensity as opposed to T17 (minor state peaks only due to overlap in $^{15}N$ for crosspeaks) and T18 peaks. For all proteins with the exception of D3-L39V, the trend observed for W43ε transitions is significantly different than that observed for T17, suggesting that different modes of exchange are experienced at each position.

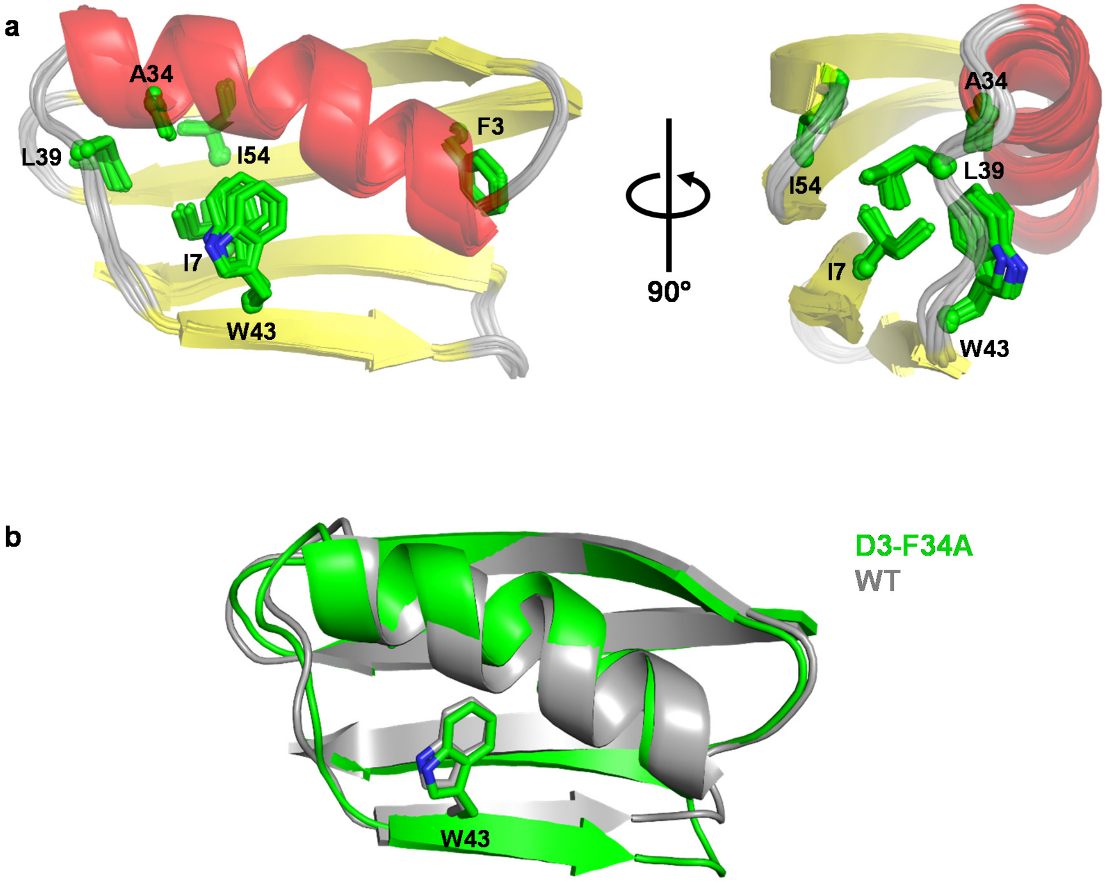


**Supplementary Figure 11. NMR solution structure of D3-F34A. a,** An NMR ensemble of 10 conformers (lowest energy) is shown for D3-F34A, with mutations from the wild type (WT) as well as W43 and A34 shown as green sticks. Structural elements, as assigned by NMR chemical shift indices,[41] are colored in red, yellow, and grey for α-helices, β-sheets, and loops respectively. **b,** An overlay of the D3-F34A structure (green) and the WT crystal structure (grey, PDB ID: 1PGA)[18] shows that their W43 side chains adopt nearly identical conformations.

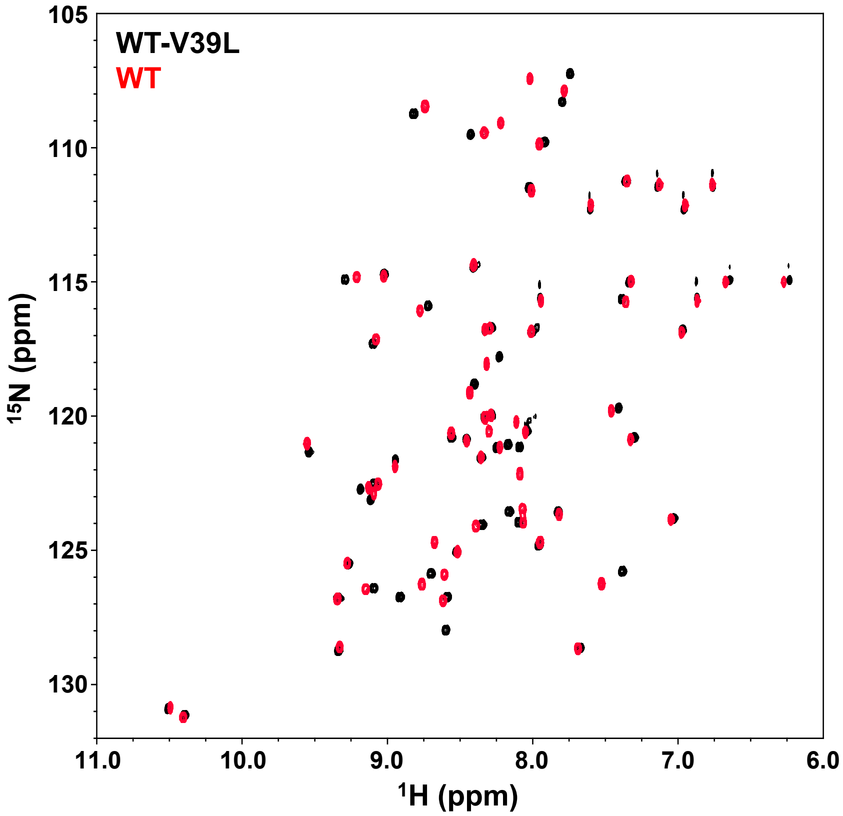


**Supplementary Figure 12. Overlay of WT-L39V and WT Gβ1 $^1$H-$^{15}$N HSQCs.** An overlay of the HSQC spectra of WT-L39V (black) and WT Gβ1 (red) shows high spectral similarity suggesting a similar structure for both proteins.

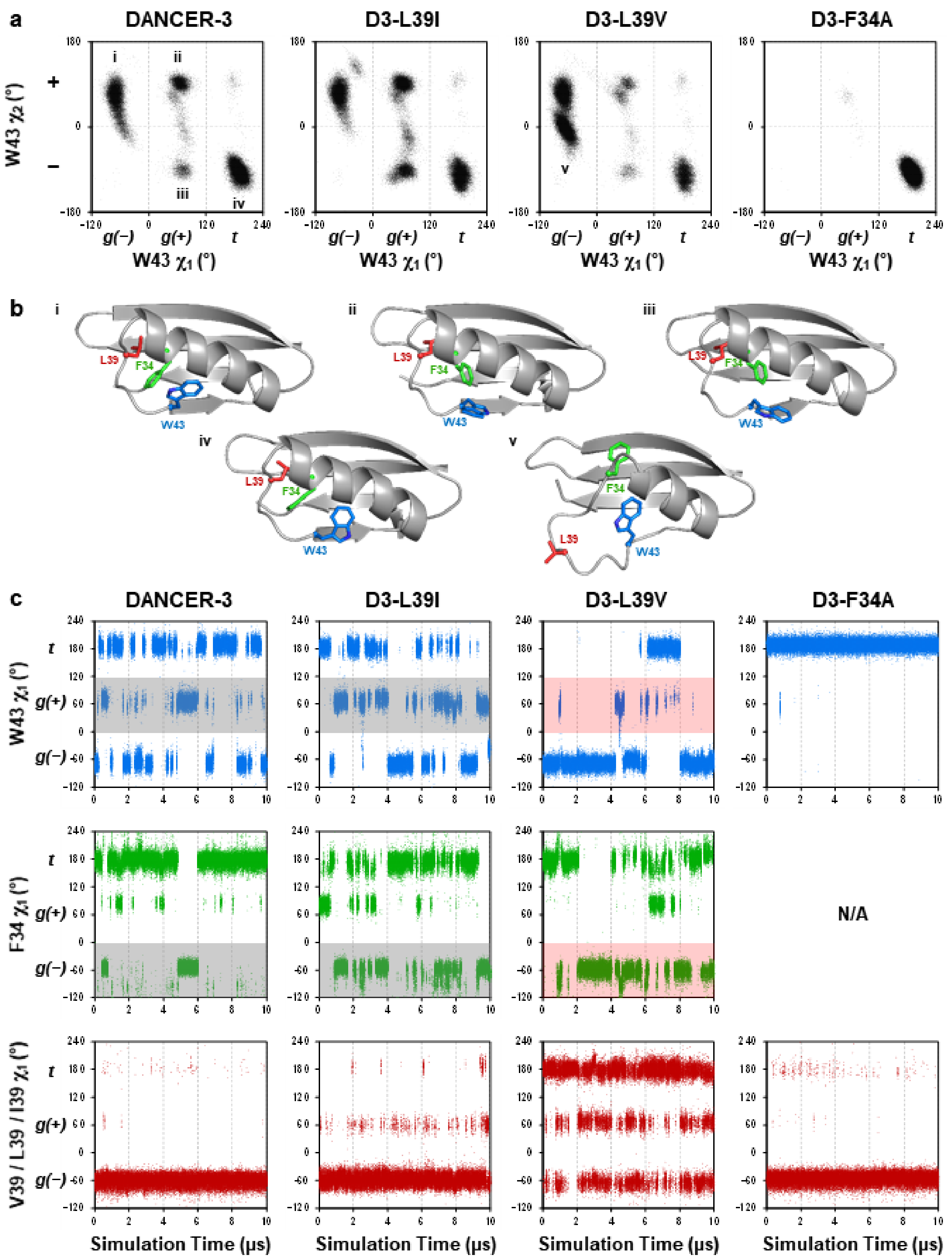


**Supplementary Figure 13. Conformational plots from molecular dynamics simulations of selected Gβ1 variants. a,** Dihedral angle plots for Trp43 χ1 and χ2 dihedral angles over the course of 10 μs of simulation time, sampled every 200 ps. **b,** Representative structures showing conformations of Trp43 (blue), Phe34 (green), and Leu39 or Val39 (red) as well as backbone conformations in each major conformational state adopted over the course of the simulations, identified in **a** using Roman numerals. **c,** $\chi_1$ dihedral angles vs time plots for Trp43, Phe34, and Leu39/Val39/Ile39 demonstrate concerted motions between Trp43 and Phe34 in DANCER-3 and D3-L39I, as shown by the simultaneous adoption of a *g(–)* conformation in Phe34 when Trp43 adopts a *g(+)* conformation (shown as grey shaded regions). These motions become uncoupled in D3-L39V (shown as red shaded regions). No concerted motions to either Trp43 or Phe34 are observed at position 39.

**Supplementary Table 1.** Dissociation constants of dimeric Gβ1 variants

| **Variant** | **Dimer $K_d$ (μM)** |
|---|---|
| WT-A34F | 27 ± 4 [a] |
| DANCER-3 | 43 ± 3 |
| D3-L39I | 16 ± 2 |
| D3-F3Y | 66 ± 5 |
| D3-I7L | 62 ± 8 |
| D3-L39V | 31 ± 3 |
| D3-I54V | 240 ± 20 |
| WT-A34F/V39L | 116 ± 5 |

[a] Value from Jee J., *et al*. (2008).

**Supplementary Table 2. Summary of NOE restraints and structural statistics [a]**

| | |
|---|---|
| **PDB ID** | 6NJF |
| **Distance Restraint Statistics** | |
| Number of NOEs | 618 |
| Short Range (\|i – j\| ≤ 1) | 336 |
| Medium Range (1 < \|i – j\| < 5) | 88 |
| Long Range (\|i – j\| ≥ 5) | 194 |
| **MolProbity Ramachandran Plot Statistics (%)** | |
| Residues in most favored regions | 96.3 |
| Residues in allowed regions | 3.7 |
| Residues in disallowed regions | 0.0 |
| **Average RMSD to mean (Å)** | |
| Backbone (mean ± 1 S.D.) | 0.29 ± 0.06 |
| Heavy Atom (mean ± 1 S.D.) | 0.79 ± 0.09 |
| **Structure Quality Factors (Raw / Z-score [b])** | |
| MolProbity clash score | 16.85 / –1.37 |
| Procheck G-factor (phi & psi) | –0.26 / –0.71 |
| Procheck G-factor (all) | –0.41 / –2.42 |
| Verify 3D | 0.42 / –0.64 |
| ProsaII (negative) | 0.56 / –0.37 |

[a] Analyzed for the 10 lowest energy structures for each designed protein using CYANA[40] and MolProbity[42]

[b] With respect to mean and standard deviation for a set of 252 X-ray structures with sequence lengths ≤ 500, resolution ≤ 1.80 Å, and R-free ≤ 0.28
